# An image analysis pipeline to quantify the spatial distribution of cell markers in stroma-rich tumors

**DOI:** 10.1101/2025.04.28.650414

**Authors:** Antoine A. Ruzette, Nina Kozlova, Kayla A Cruz, Taru Muranen, Simon F. Nørrelykke

## Abstract

1.

Aggressive cancers, such as pancreatic ductal adenocarcinoma (PDAC), are often characterized by a complex and desmoplastic tumor microenvironment rich in stroma, a supportive connective tissue composed primarily of extracellular matrix (ECM) and non-cancerous cells. Desmoplasia, which is a dense deposition of stroma, is a major reason for therapy resistance, acting both as a physical barrier that interferes with drug penetration and as a supportive niche that protects cancer cells through diverse mechanisms. A precise understanding of spatial cell interactions within the tumor microenvironment in stroma-rich cancers is essential for optimizing therapeutic responses. It allows detailed mapping of stromal-tumor interfaces, comprehensive phenotyping of diverse cell types and their functional states, and insights into changes in cellular distribution and tissue architecture, thus leading to an improved assessment of drug responses. Recent advances in multiplexed immunofluorescence imaging have enabled the acquisition of large batches of whole-slide tumor images, but scalable and reproducible methods to analyze the spatial distribution of cell states relative to stromal regions remain limited. To address this gap, we developed an open-source computational pipeline that integrates QuPath (Bankhead et al. 2017), StarDist (Schmidt et al. 2018), and custom Python scripts to quantify biomarker expression at a single- and sub-cellular resolution across entire tumor sections. Our workflow includes: (i) automated nuclei segmentation using StarDist, (ii) machine learning-based cell classification using multiplexed marker expression, (iii) modeling of stromal regions based on fibronectin staining, (iv) sensitivity analyses on classification thresholds to ensure robustness across heterogeneous datasets, and (v) distance-based quantification of the proximity of each cell to the stromal border. To improve consistency across slides with variable staining intensities, we introduce a statistical strategy that translates classification thresholds by propagating a chosen reference percentile across the distribution of marker-related cell measurement in each image. We apply this approach to quantify spatial patterns of distribution of the phosphorylated form of the N-Myc downregulated gene 1 (NDRG1), a novel DNA repair protein that conveys signals from the ECM to the nucleus to maintain replication fork homeostasis, and a known cell proliferation marker Ki67 in fibronectin-defined stromal regions in PDAC xenografts. The pipeline is applicable for the analysis of various stroma-rich tissues and is publicly available: https://github.com/HMS-IAC/stroma-spatial-analysis-web.

**Summary paragraph:** Our study introduces a scalable and reproducible image analysis pipeline that quantifies spatial biomarker distributions relative to the stroma in tumor tissues using open-source tools. By modeling cell-level intensity distributions and calibrating classification thresholds across heterogeneous images, we uncover spatially organized patterns of stroma sensing, DNA damage, and proliferative response in pancreatic tumors. This approach enables robust, quantitative analysis of tumor-stroma interactions and is readily adaptable to other tumor types and biomarker panels, providing a valuable resource for spatial pathology and tumor microenvironment research.

## 3. Introduction

Tumor microenvironment (TME) is a complex neighborhood that plays an active role in tumor progression, metastasis, immune evasion, and influences the efficacy of anti-cancer agents (Valkenburg, de Groot, and Pienta 2018; Xu et al. 2022). TME consists of cancer cells, cancer-associated fibroblasts (CAFs), immune cells, endothelial cells, pericytes, neuronal cells, and adipocytes. Central to the TME is the stroma, a supportive framework composed primarily of connective tissue, extracellular matrix (ECM) proteins, blood vessels, fibroblasts, and immune cells. CAFs are known to secrete excessive amounts of ECM proteins in desmoplastic cancers, thus regulating the stromal density within a tumor. Desmoplasia is a hallmark of pancreatic ductal adenocarcinoma (PDAC), a malignancy marked by aggressive behavior, poor prognosis, and a dense, fibrotic stroma that limits drug penetration, fosters an immunosuppressive environment, and orchestrates signaling events leading to therapy resistance (Kozlova et al. 2020; Ho, Jaffee, and Zheng 2020; Halbrook et al. 2023). Precise understanding of the spatial organization of the TME is essential for decoding the complexity of cell-stroma interactions, and for predicting treatment efficacies.

Each cell population within TME can be defined by immunostaining with specific markers (Hu et al. 2023). While epithelial cells can be distinguished by positive staining of various cytokeratins (Karantza 2011), the visualization of stromal compartment within a tumor can be done via Masson’s Trichrome stain that stains for collagen and fibrin for brightfield imaging (Masson 1929), or by immunofluorescent staining of matrix proteins such as collagens, fibronectin and laminins (Kozlova et al. 2020). Recent advances in multiplexed immunofluorescence imaging allow for high-resolution visualization of multiple biomarkers across entire tumor sections. In parallel, tools to analyze bioimages have emerged (Cimini et al. 2024). Several software platforms support general microscopy image analysis—including open-source tools such as QuPath (Bankhead et al. 2017), CellProfiler (Stirling, Carpenter, and Cimini 2021; Stirling et al. 2021), and Fiji (Schindelin et al. 2012; Schneider, Rasband, and Eliceiri 2012; Schindelin et al. 2015; Rueden et al. 2017), and commercial packages like HALO (Indica Labs, Albuquerque, NM, USA) and Visiopharm (Hørsholm, Denmark). While each offers strengths, most fall short in modeling spatial relationships between cell phenotypes and stromal structures across heterogeneous, multi-channel datasets. Commercial solutions provide turnkey pipelines but lack flexibility for custom analysis; open-source tools often require manual threshold tuning or lack built-in support for stromal modeling and spatial distance quantification. Moreover, ensuring reproducibility in the life sciences is crucial, and the price barrier of commercial solutions hampers accessibility and reproducibility. Open-source workflows offer greater transparency but require a careful attention to understand the sensitivity of methods used under image variability (Miura and Nørrelykke 2021).

To address these challenges, we developed a robust, open-source pipeline for quantifying the spatial distribution of cell markers relative to stromal borders in tumor tissues. Built on QuPath (Bankhead et al. 2017), our workflow integrates nuclei segmentation with StarDist (Schmidt et al. 2018), machine learning-based cell classification at cellular and sub-cellular resolution, stromal region modeling, sensitivity analysis, and spatial distance measurements. We also introduce a statistical mapping method that translates intensity-based thresholds across heterogeneous images using percentile propagation in continuous distributions of cell measurements. The pipeline leverages Python-based tools for data aggregation and spatial visualization, facilitating seamless analysis of large, multi-image datasets.

Our previous study identified N-Myc downregulated gene 1 (NDRG1) as a novel DNA repair protein able to convey signals from the ECM to the nucleus to maintain replication fork homeostasis and mediate resistance to chemotherapies. In this work we present the pipeline ‘in action’ on multiplexed immunofluorescence images of PDAC xenografts treated with chemotherapy. This pipeline defines spatial distribution of cells expressing pan-cytokeratin (PDAC cells), phosphorylated NDRG1 (a stromal sensor), and Ki67 (a marker of cell proliferation) in relation to stromal border modeled by fibronectin staining. Our analysis reveals the enrichment of the phospho-NDRG1- and Ki67-positive cells at the stromal border and their detailed spatial distribution based on signal intensity. The pipeline is scalable, robust to staining variability, and adaptable to other biomarker panels, offering a reproducible framework for spatial analysis of components of the tumor microenvironment in relation to stromal regions and their border.

## 4. Methods

### 4.1. Xenograft studies, tissue processing and image acquisition

Pancreatic adenocarcinoma cell line AsPC was a kind gift from Dr. Nada Kalaany (Boston Children’s Hospital, Boston, MA). All animal studies were performed according to protocols approved by the Institutional Animal Care and Use Committee at Beth Israel Deaconess Medical Center (BIDMC). Athymic male nude mice (NU/J) were purchased from Jackson labs (#002019). AsPC tumor cells (1×10^6^) were injected subcutaneously in one flank per mouse in 1:1 mix of Matrigel and PBS when the mice were 8-12 weeks old. Tumor growth was monitored thrice weekly. Once the tumors reached 200 mm^3^ in volume, animals were treated with gemcitabine 25 mg/kg twice per week. Entire cohort of animals was sacrificed when tumor size of at least one animal was close to reach 1000 mm^3^.

Xenograft tumor tissues were processed, paraffin embedded and cut, deparaffinized in xylene and rehydrated in a descending ethanol series. Deparaffinized sections were subjected to antigen unmasking in SignalStain® Citrate Unmasking Solution (CST #14746) followed by permebealization in 1% Triton X-100, and blocking of nonspecific binding in TBST/5% normal goat serum solution. Sections were later stained with mouse anti-Fibronectin (ab6328-250, Abcam), Pan Cytokeratin (AE1/AE3), Alexa Fluor™ 488 conjugated (53-9003-82, eBioscience) and rabbit phospho-NDRG1Thr 346 (#5482S, CST) or Ki67 (D3B5) Rabbit mAb (Alexa Fluor® 647 Conjugate) (#12075, CST). Appropriate secondary antibodies were used and whole section images were acquired using Olympus BX-UCB whole slide scanner equipped with Olympus UPLSAPO 20x and VS-ASW v2.7 software, tile registration was performed using the scanner’s proprietary software. Each image consisted of four fluorescence channels: DAPI (nuclei), TRITC (pan cytokeratin), FITC (fibronectin), and CY5, labeling either phosphorylated NDRG1 (dataset 1) or Ki67 (dataset 2). Each field covered approximately 10,000 *µ*m × 10,000 *µ*m (30,000 × 30,000 pixels at 0.3215 *µ*m per pixel) and contained roughly 100,000 cells per image. Dataset 1 (pNDRG1) contained a total of 496,936 cells across all images; dataset 2 (Ki67) contained 454,556 cells. The methods described below are channel-agnostic and serve as a generalizable pipeline for multi-marker spatial analysis in tissue sections.

### 4.2. Modeling the stromal region and its border

The stroma in PDAC is composed of a complex mixture of cells and extracellular matrix (ECM) components, which makes delineation of precise stromal borders nontrivial. Fibronectin, a key ECM protein enriched in the stromal compartment, was used as a surrogate marker to define stromal regions. To reduce noise and imaging artifacts, fibronectin channel intensities were first smoothed using a Gaussian filter; the sigma parameter was tuned empirically based on expert input to balance edge preservation with noise reduction and then confirmed with a sensitivity analysis. A threshold-based pixel classifier was then applied to segment stromal versus non-stromal tissue. Pixels exceeding the fibronectin intensity threshold were classified as stromal. This threshold was selected by an experienced cancer biologist and later translated across the dataset using a percentile-based statistical propagation method (**Box 1**). The border of the resulting binary mask was used as the stromal edge and served as the spatial reference for downstream analyses (**Figure 1A**).

**Figure 1:**
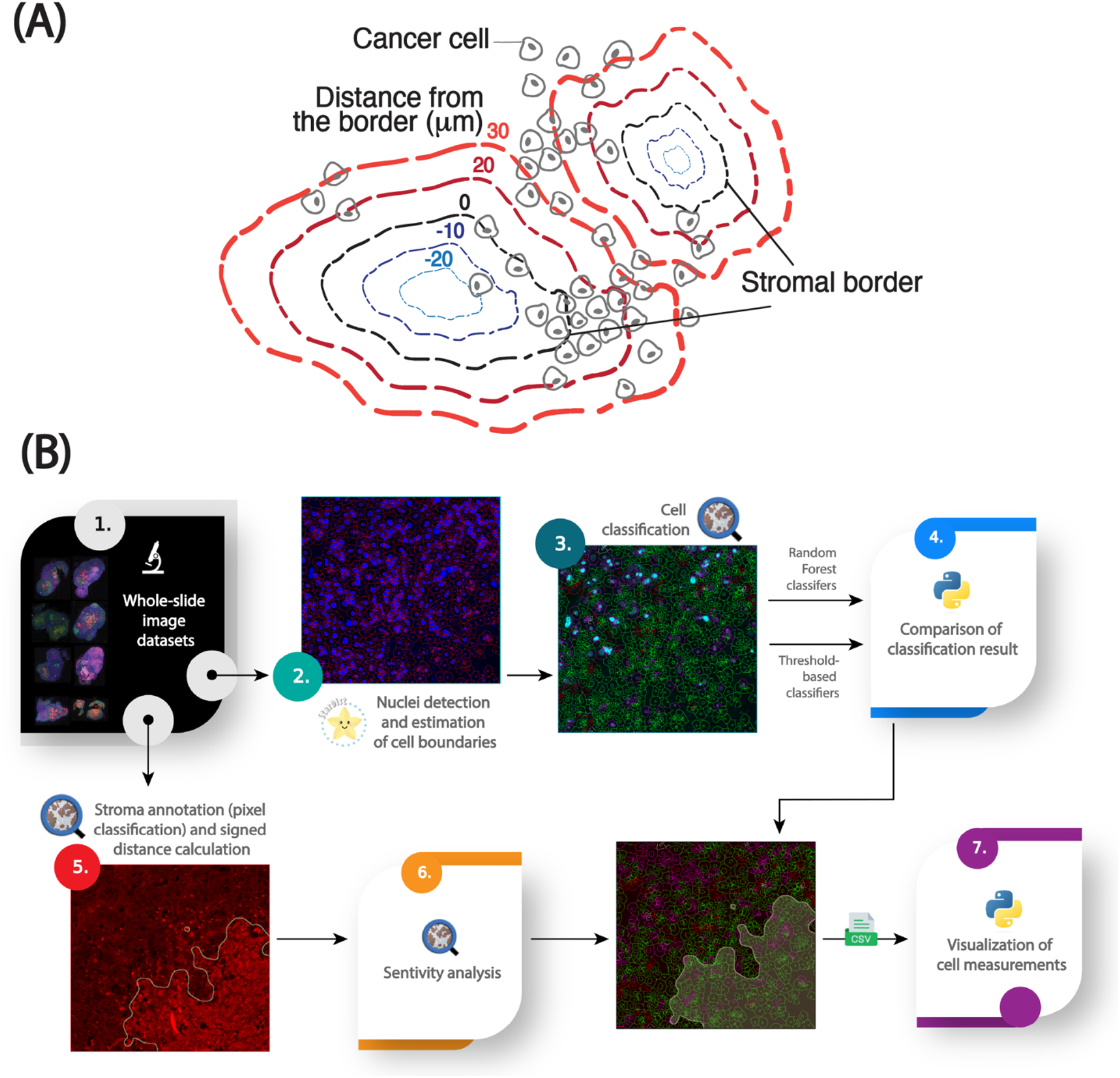
An image analysis pipeline for quantifying the spatial distribution of cellular markers in stroma-rich tumors (adapted from (Kozlova et al. 2025)). **(A)** Graphical abstract of the spatial model for stroma-dense tissues. **(B)** Workflow for analyzing cellular and subcellular fluorescent markers relative to a modeled stromal border, including nuclei segmentation with StarDist, cell boundary estimation using QuPath, Random Forest models for cell classification, pixel threshold-based stroma annotation, 2D signed distance calculations for spatial analysis, and data visualization in Python.

**Figure 2:**
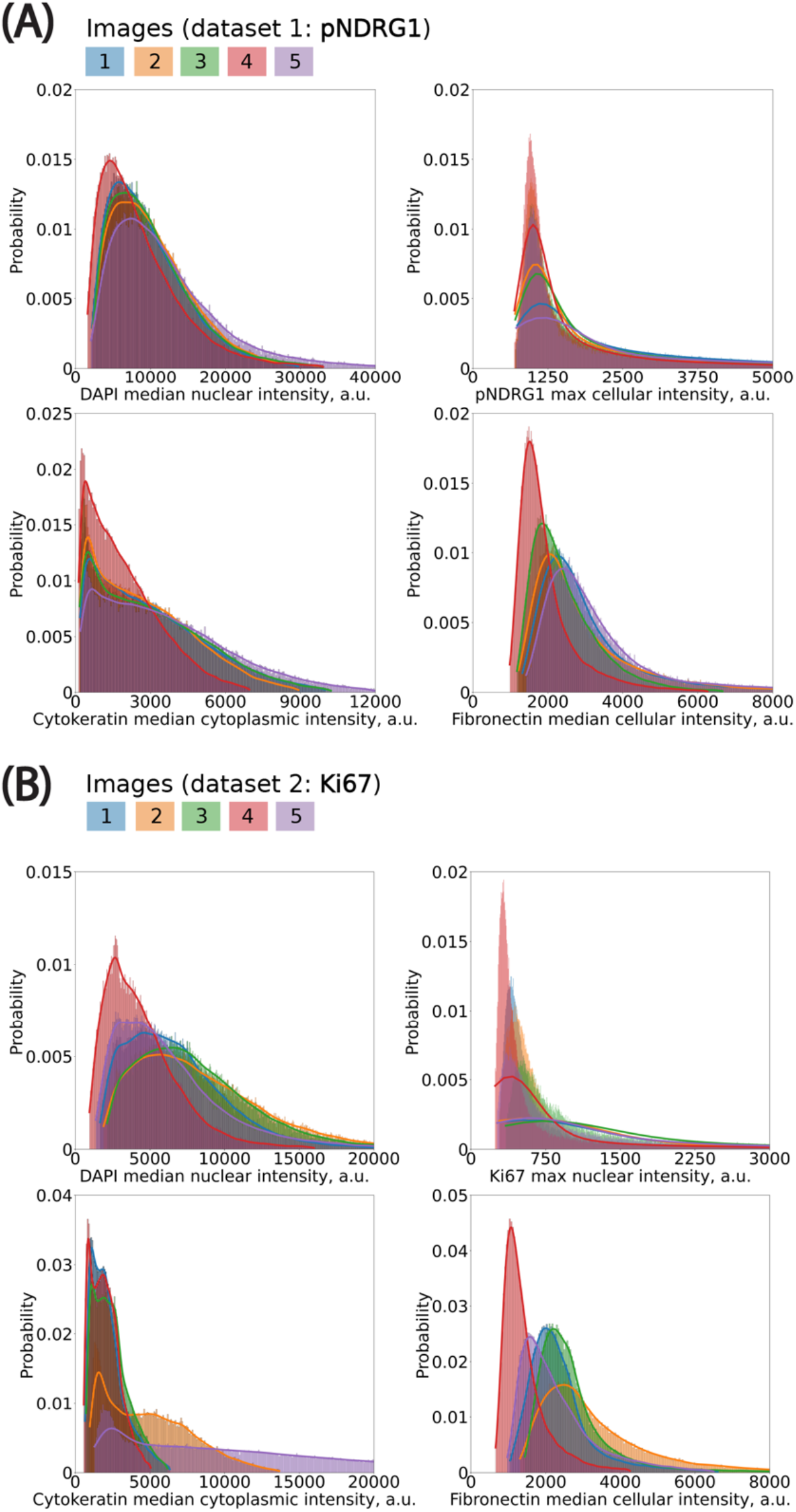
Histograms in cell measurements of interest. **(A)** Dataset contains 5 images. Bin size (in *µ*m): DAPI, 150; pNDRG1, 5; KER, 50; FN, 20.**(B)** Dataset contains 5 images. Bin size (in *µ*m): DAPI, 50; Ki67, 5; KER, 75; FN, 40. y-axis represents the percentage of total observations that fall into each bin.

### 4.3. Image analysis components

#### Nuclei detection and cell boundary approximation

Nuclei were segmented using the pre-trained StarDist model (“dsb2018_heavy_augment.pb”, available at QuPath models repository: https://github.com/qupath/models/tree/main/stardist) via the StarDist extension in QuPath (v0.5). StarDist models nuclei as star-convex polygons and has demonstrated high accuracy across diverse nuclear morphologies (Schmidt et al., 2018). The output was a set of nuclear regions of interest (ROIs), each representing a single cell nucleus. To approximate whole-cell boundaries, QuPath’s built-in cytoplasmic expansion algorithm was applied, radially expanding each nucleus by 5 *µ*m. Cells were filtered based on nuclear area to exclude likely artifacts: nuclei smaller than the 5th percentile or larger than the 99th percentile were discarded, as they likely represented immune cells, debris, or fused nuclei caused by tissue processing artifacts.

#### Distributions of cell intensity measurements

For each cell, fluorescence intensity features were extracted from the relevant subcellular compartments (nucleus, cytoplasm, whole cell) in QuPath. Signal intensity distributions were visualized and compared across images in the same batch using fixed bin widths. To reduce the influence of outliers, intensities were clipped at the 1st and 99th percentiles. No normalization was applied, preserving the raw distribution.

#### Cell classification

Cells were classified as marker-positive or -negative using either a supervised classifier or a percentile-based intensity thresholding approach:

##### 1. Supervised classification

A Random Forest classifier was trained within QuPath using a manually annotated subset of cells. The training set focused on intensity-based features relevant to the marker being evaluated (e.g., maximum nuclear intensity for Ki67 foci). Only intensity-related features were used to simplify model interpretation and reproducibility.

##### 2. Percentile-based thresholding

A fixed intensity percentile was selected from the cell population within a reference image and applied across all images. This thresholding approach classified the top N% of cells by marker intensity as “positive,” enabling harmonized comparisons across heterogeneous images.

Confusion matrices and derived standard metrics (agreement percentage) were computed to evaluate agreement between these two classification strategies. High agreement between the methods indicated classifier stability, while deviations flagged potential staining inconsistencies or classifier bias.

#### Percentile mapping of expert-chosen classification thresholds

To propagate expert-defined intensity thresholds across images with varying intensity distributions, we implemented a statistical mapping method based on percentile preservation. For each image, the distribution of intensity values for a given marker was modeled using the best-fit probability distribution (minimizing least-squares error). The expert-selected intensity threshold in a reference image was then mapped to the corresponding percentile, and this percentile was used to determine equivalent thresholds in other images. The algorithm is statistically described in **Box 1**. The resulting mapped thresholds were compared to machine learning-based classifications. Using one method’s labels as “ground truth,” confusion matrices were computed to evaluate accuracy and agreement. This percentile mapping approach enabled reproducible thresholding across large heterogeneous datasets while preserving expert intent.

#### 2D signed distance between cells and the stromal border

We used the signed Euclidean distance between each cell’s nuclear centroid and the nearest point on the stromal border to quantify spatial relationships between tumor cells and the stroma. Positive distances indicate cells located outside, while negative distances represent cells within the fibronectin-defined stromal region. Cells located at the interface thus have distances close to 0 *µ*m. Cells were binned by distance using 10 *µ*m intervals, approximately corresponding to one cell diameter. These distance bins formed the basis for downstream spatial analyses and visualization of intensity over distance from the stroma.

#### Sensitivity analysis on intensity thresholds

To assess the robustness of the stromal mask and downstream spatial results, we conducted sensitivity analyses on two key parameters: the Gaussian smoothing sigma and the fibronectin intensity threshold. For each parameter combination, the stromal mask was recomputed, followed by recalculation of the signed distances and intensity-distance correlations.

As a summary metric, we calculated the Pearson correlation coefficient between marker intensity *I* and distance to the stromal border *d*, separately for cells inside and outside the stroma. The correlation coefficient *r* was computed as:

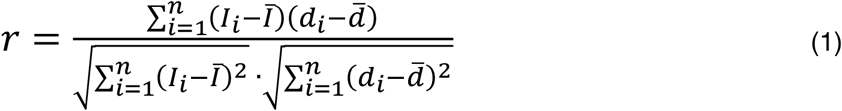

where:

- *I*_*i*_ is the signal intensity for the *i*-th cell
- *d*_*i*_ is the signed distance to the stromal border for the *i*-th cell
- *Ī* is the mean signal intensity
- 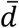 is the mean signed distance

For each image and parameter setting, correlation coefficients were computed separately for regions before and after the zero-distance boundary. Because both correlation values were derived from the same image under identical conditions, the data were treated as paired measurements. Differences in correlation values across the two distance partitions (inside and outside the stroma) were assessed using a two-sided Wilcoxon signed-rank test, a non-parametric test for paired data. Mean correlation values and standard errors were estimated via 500 bootstrap iterations across images.

#### Visualization

Final cell-level data, including marker intensities, class labels, and signed distances, were exported from QuPath and analyzed in Python 3.11. Cells were grouped into 10 *µ*m distance bins, and average marker intensity with bootstrapped standard error was computed for each bin. Profiles were generated for marker-positive and marker-negative populations separately. To assess differences in spatial patterns, bin-wise subtractions were performed between groups (e.g., Ki67-positive vs. Ki67-negative cancer cells). The standard error of the difference in bin means was computed using error propagation:

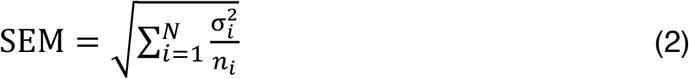

where:

- SEM is the propagated standard error
- *σ*_*i*_ is the standard deviation of signal intensities in a cell population, and *n*_*i*_ the number of cells in the population

Spatial distributions were visualized as line plots with error bands, highlighting marker-specific spatial gradients in relation to the stromal border (**Figure 5**).

## 5. Results

### 5.1. Batch variability and statistical propagation of thresholds

To ensure the reliability of downstream analyses and aggregation across images, we first assessed the consistency of per-cell fluorescence intensity distributions within each image batch. While some variation is expected due to acquisition conditions and biological heterogeneity, we observed overall alignment in signal profiles across both datasets (**Figure 3**). Most notably, the distribution of median nuclear intensity in the DAPI channel remained highly consistent across all five images, indicating uniform nuclear staining and robust segmentation quality (**Figure 3A**). Although Image #5 showed the greatest deviation, its values remained within acceptable bounds and did not compromise downstream analysis. In contrast, the pan-cytokeratin (TRITC) channel—used as a marker to distinguish cancer cells from non-cancer cells—exhibited more substantial variability, particularly in median cytoplasmic signal intensity per cell (**Figure 3B**). Two of the five images showed significantly elevated pan-cytokeratin intensities. Rather than exclude these images, we retained them in order to test the robustness of our threshold calibration strategy under conditions of variable staining. This variability offered an opportunity to assess how well our pipeline handles intensity heterogeneity without compromising classification consistency.

**Figure 3:**
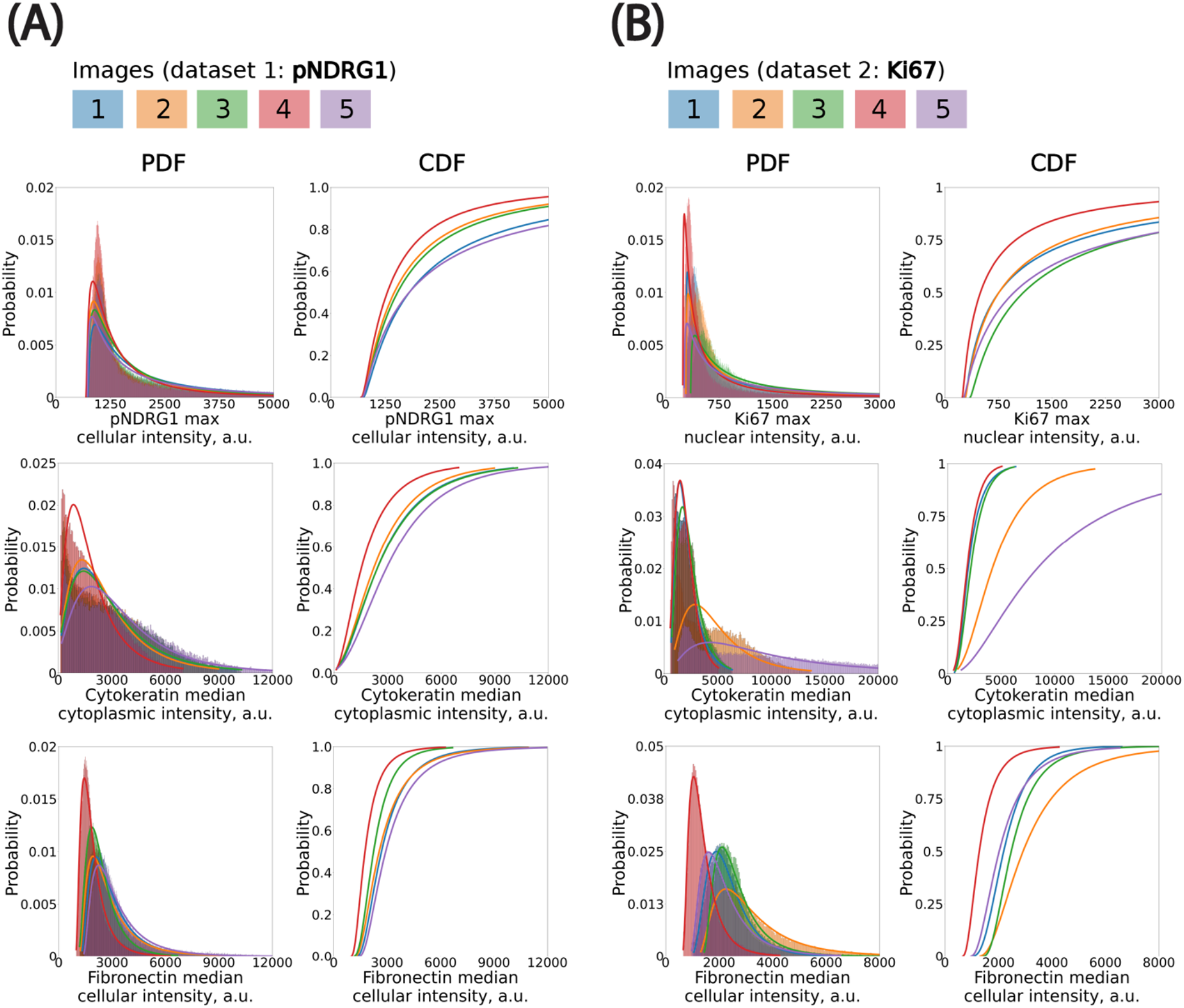
Fitting probability distributions to histograms to propagate classification thresholds using percentile mapping across images. **(A)** pNDRG1 dataset. Bin size (in *µ*m): DAPI, 150; pNDRG1, 5; KER, 50; FN, 20. **(B)** Ki67 dataset. Bin size (in *µ*m): DAPI, 50; Ki67, 5; KER, 75; FN, 40. All x-axis are clipped to highlight the left elbow of distributions. CDF: Cumulative Distribution Function. PDF: Probability Density Function.

We modeled the cell-level fluorescence histograms in each image using continuous, positive-valued probability distributions (**Box 1**). We evaluated multiple candidate distributions and found that the log-normal distribution consistently provided the best fit, as assessed by least-squares error. This distribution was especially well-suited to the skewed, elbow-shaped profiles commonly observed in fluorescence intensity histograms across both cytoplasmic and nuclear compartments. While fluorescence signals originate from discrete photons and are digitized into grey levels, the continuous log-normal distribution remains a good approximation at high photon counts—conditions under which the discreteness of the signal becomes negligible due to the Central Limit Theorem. Accordingly, we used log-normal fits for all channels in subsequent statistical threshold propagation.

Expert-defined thresholds were established in a single reference image for each marker (excluding DAPI), and these values were mapped to the remaining images by preserving their percentile rank within the fitted log-normal distribution. This ensured that a consistent fraction of cells—corresponding to the top X% by intensity—was classified as marker-positive in each image, adapting the absolute threshold to the local intensity distribution. We applied this method to both phospho-NDRG1 and Ki67 image datasets. This percentile-based propagation approach offered advantages over using a fixed global threshold or a threshold specifically picked for each image. In practice, it improved the selection of marker-positive cells by statistically accounting for image-specific differences in staining intensity. It also provided a way to only have to manually select one threshold. We expect this method to be even more critical in studies with larger batch effects or multi-site imaging datasets, where inter-image variability is more pronounced.

### 5.2. Application case 1: Revealing the spatial distribution between the phosphorylated form of NDRG1, a novel DNA repair protein required for stroma-induced chemoresistance, and stromal border

To demonstrate the utility of our pipeline in uncovering biologically meaningful spatial patterns, we analyzed the distribution of cells expressing high levels of phosphorylated NDRG1 (pNDRG1), a novel DNA repair protein implicated in chemoresistance and replication fork stability, in PDAC AsPC xenografts treated with gemcitabine, a standard-of-care chemotherapy used in the treatment of pancreatic cancer. Our previous work showed that phosphorylation of NDRG1 is regulated by ECM and adhesion receptors leading to the enrichment of pNDRG1-positive cells near tumor-stroma interfaces in PDAC SW1990 xenografts (Kozlova et al. 2025). Given that NDRG1 is localized both in the cytoplasm and the nucleus, we used maximum whole-cell intensity as the primary feature for classification. Additional features included median cytoplasmic pan-cytokeratin intensity (to select for cancer cells) and a median cellular fibronectin intensity (for stromal modeling). Fibronectin is a secreted matrix deposited outside of the cells; the median cellular intensity was only used as a proxy to visualize its distribution in a similar fashion (at the cell level) than the other markers.

We trained a Random Forest classifier (QuPath, built-in) on a balanced set of 318 manually labeled cells spanning four classes: pan-cytokeratin-positive, pan-cytokeratin-negative, pNDRG1-positive, and pNDRG1-negative (**Figure 4A**). In parallel, we designed a threshold-based classifier using percentile-propagated cutoffs from a reference image (Image #1), as described in **Box 1**. Agreement between the machine learning and threshold-based methods was high (87.7%, **Figure 4C**), confirming the consistency of classification across heterogeneous images. When comparing the percentile-propagation method to a fixed global threshold, we observed an average increase in classification agreement of 1.36% (±6.88% SD, **Table S7**). These findings highlight that the necessity of accounting for intensity distribution variations is highly context dependent. In some cases, a single-threshold approach can approximate the discriminative power of a trained classifier, whereas in others, statistical propagation may be more appropriate. As a matter of illustration, when applying the set of thresholds of image #4 on the rest of the batch, the agreement drops to 78.7%. In other words, using statistical propagation averages out the variations observed when using a single set of thresholds for all images–providing a more robust way of picking a single set of thresholds for a whole batch. Full classification parameters and translated thresholds are provided in the supplementary material (**Tables S1–S3**).

**Figure 4:**
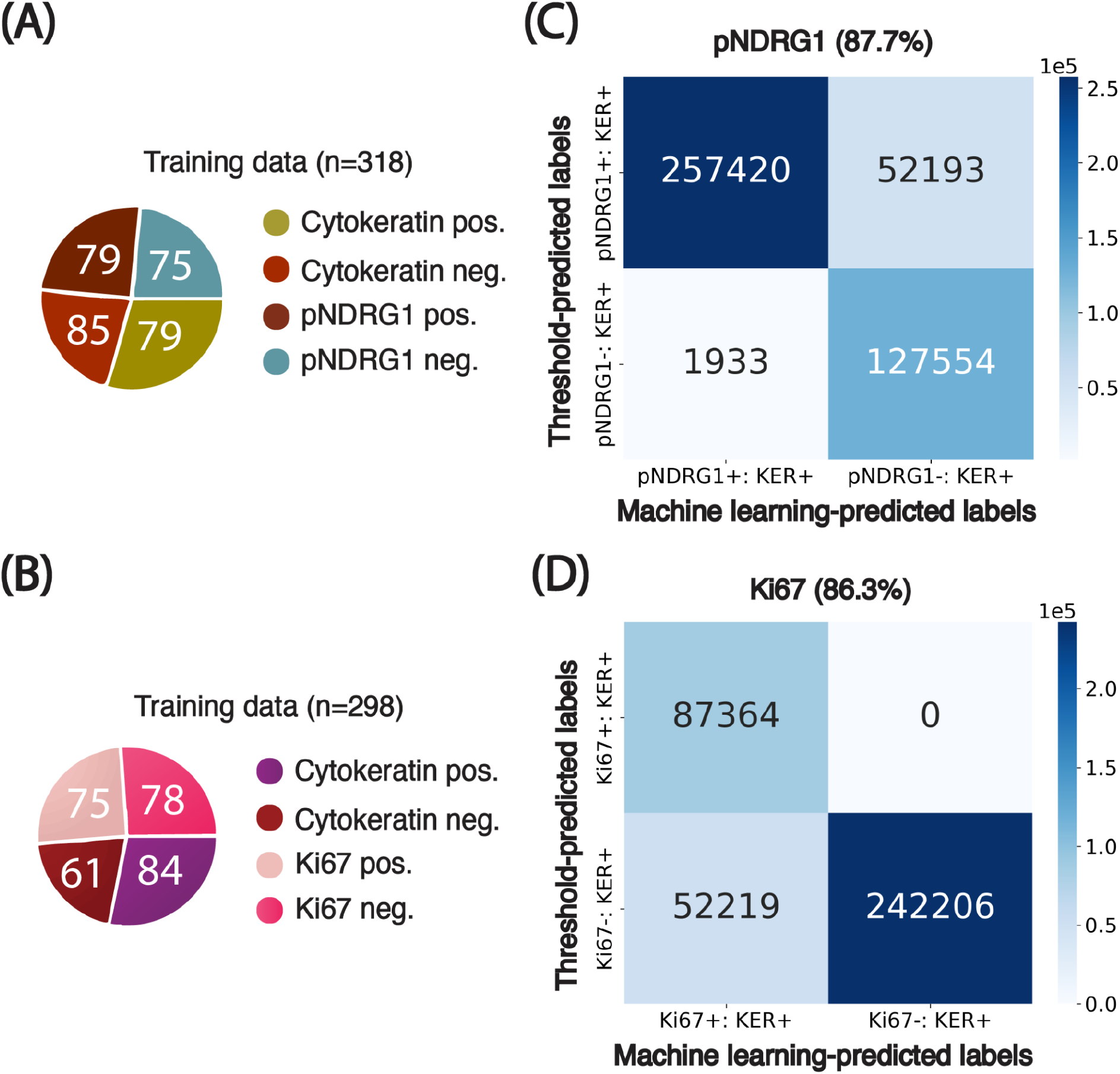
Agreement of results from machine learning-based and threshold-based classifiers. **(A)-(B)** Breakdown of the training set curated for cells in both pNDRG1 and Ki67 images. Numbers inserted in charts indicate the number of training examples per class. **(B)-(C)** Confusion matrices comparing threshold-based and machine learning-based classification of cells based on cellular marker signal intensity. Percentages indicate the agreement between classification results from the two methods. **KER:** Pan-cytokeratin.

**Figure 5:**
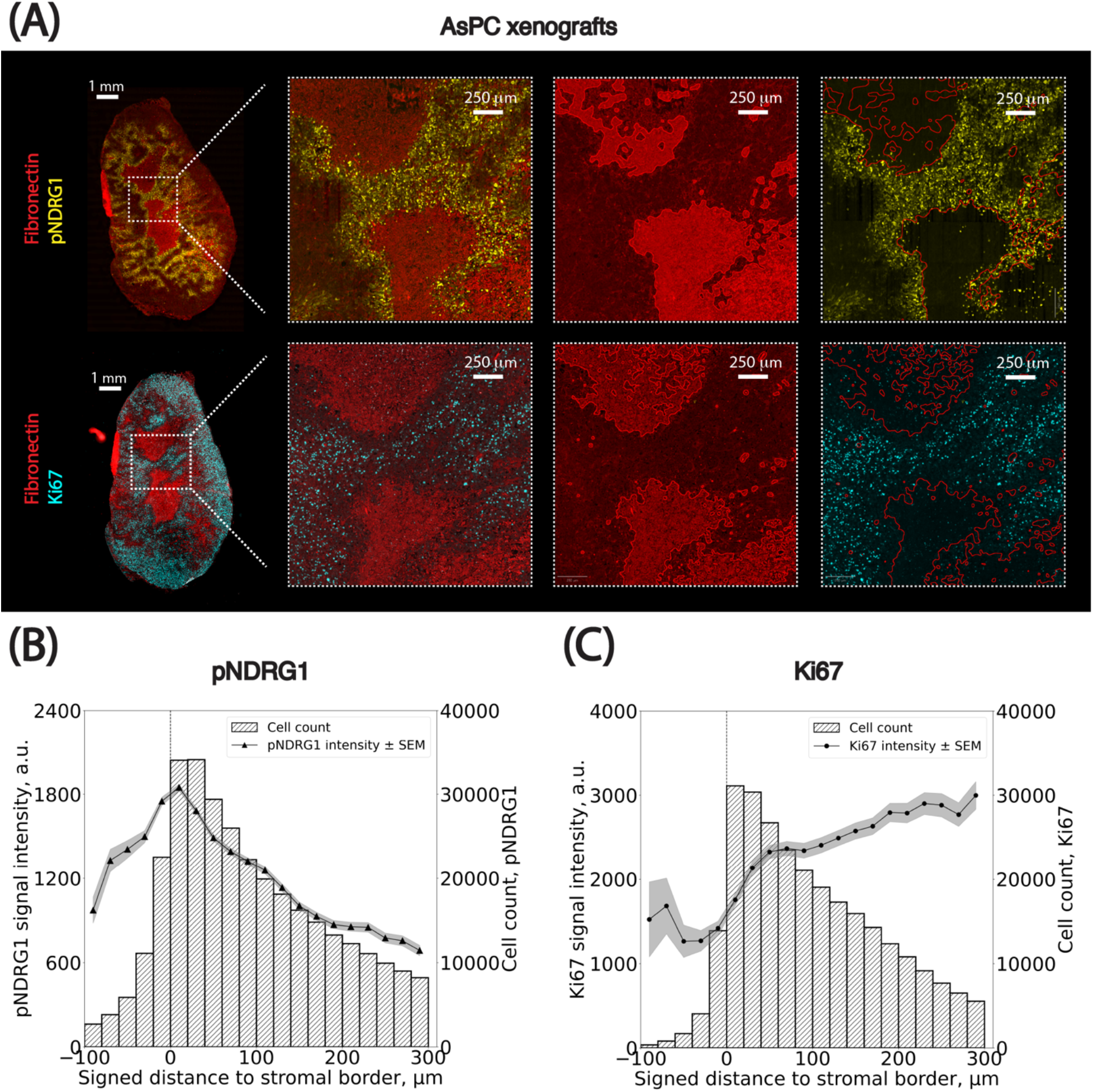
Spatial distributions of pNDRG1- and Ki67-positive cancer cells relative to the stromal border. **(A)** Representative images of the PDAC AsPC xenograft sections stained for Fibronectin (red), phospho-NDRG1 (yellow) and Ki67 (cyan) used for spatial analysis. **(B-C)** Spatial analysis of pNDRG1 and Ki67 signal intensity in cancer cells as a function of their signed distance to the closest stromal border (x-axis, in *µ*m). Cells positive for pNDRG1 or Ki67 were grouped into bins based on their distance from the stromal border. The left y-axis shows the average maximum cellular signal intensity for pNDRG1- or Ki67-positive cells (black dots) per bin, with standard error of the mean (SEM) indicated as grey overlays. The right y-axis represents the number of cells per bin. Negative distances correspond to cells within stromal regions, while positive distances indicate cells outside these regions. Bin size: 10 *µ*m. The x-axis is clipped between -100 and 300 *µ*m around the stromal border. Results from the cell classification using adaptive thresholds were used.

Spatial analysis applied to AsPC xenograft images revealed that pNDRG1-positive cells were enriched near stromal borders. Visual inspection of whole-slide images showed areas of high pNDRG1 intensity in tumor regions adjacent to stroma (**Figure 5A**). Both classification methods consistently quantified this pattern. Two key trends emerged (**Figure 5B**):

1. pNDRG1 intensity peaked at the stromal borders of gemcitabine treated tumors.
2. The number of pNDRG1-positive cells within the stomal regions is much lower compared to the one outside the stromal compartment, as can be assessed by the number of cells in each bin.

These findings are in line with a general observation, that pancreatic stroma presents as a desmoplastic matrix limiting cell infiltration. The cells adjacent to the stromal border respond to mechanical sensing of the ECM which is reflected in the phosphorylation of the NDRG1. These findings also emphasize the value of incorporating intensity calibration into cross-image analyses.

### 5.3. Application case 2: The case of Ki67 revealing the spatial distribution between cell proliferation and stromal border

We next applied the pipeline to Ki67, a canonical nuclear marker of cellular proliferation, to investigate whether spatial gradients of proliferative activity exist in relation to the stromal border. Since Ki67 localization is strictly nuclear, we used maximum nuclear intensity as the key classification feature.

A Random Forest classifier was trained on 298 manually annotated cells across four classes (pan-cytokeratin-positive, pan-cytokeratin-negative, Ki67-positive, and Ki67-negative; **Figure 4C**). We also defined percentile-based thresholds from Image #1 and propagated them across all images using our statistical method (**Box 1**). Both classification approaches produced similar results, with an average agreement of 86.3% (**Figure 4D**). However, unlike pNDRG1, Ki67 classification proved more sensitive to intensity heterogeneity: using a single global threshold led to an average agreement drop of 2.9% (±10.1% SD, **Table S8**), highlighting the necessity of threshold calibration when analyzing markers with higher inter-image variability. Full classification parameters and translated thresholds are provided in the supplementary material (**Tables S4–S6**).

Spatial mapping of Ki67-positive cells revealed a clear trend: the number of actively proliferating cells was lowest within stromal regions and progressively increased with distance from the stroma-tumor interface, plateauing at approximately 300 *µ*m (**Figure 5C**). Two observations support this:

1. Ki67 signal intensity increased monotonically from 0 to ∼300 *µ*m away from the stromal border, after which it stabilized (**Figure S2**).
2. Similarly, the number of Ki67-positive cells within the stomal regions was much lower compared to the one outside the stromal compartment, assessed by the number of cells in each bin (**Figure S2**).

These findings support the notion that dense stroma creates a mechanical and a biochemical barrier that manifests in diminished exposure of cancer cells to chemotherapy, thus having a major impact on cancer cell proliferation. They also demonstrate the importance of adapting classification strategies to accommodate image-specific intensity profiles.

### 5.4. Sensitivity analysis of thresholds on the spatial distribution

We conducted a sensitivity analysis to assess the robustness of spatial trends with respect to two key parameters involved in stromal mask generation: (1) the standard deviation (σ) of the Gaussian kernel applied to the fibronectin channel and (2) the pixel intensity threshold used to distinguish stromal from non-stromal regions. We performed a grid search over both parameters and, for each combination, recalculated the stromal mask, cell-to-stroma distances, and the Pearson correlation between marker intensity and stromal proximity. Across this parameter space, spatial trends remained consistent for both markers (**Figure 6**). For pNDRG1, the correlation between intensity and distance ranged from -0.01 to 0.03 for cells inside the stroma and from -0.16 to -0.15 for cells outside the stroma. For Ki67, the correlation ranged from 0.04 to 0.07 inside the stroma and from 0.01 to 0.03 outside the stroma. These small fluctuations indicate that the observed spatial patterns are robust to reasonable variations in smoothing and thresholding. Taken together, the results confirm that the observed spatial patterns are not artifacts of specific parameter choices. This step also led us to empirically select σ = 15 for pNDRG1 and σ = 10 for Ki67, based on the parameter set that maximized smoothness of the stromal border while preserving local spatial structure. These values produced segmentation masks that were both biologically plausible and numerically stable across images.

**Figure 6:**
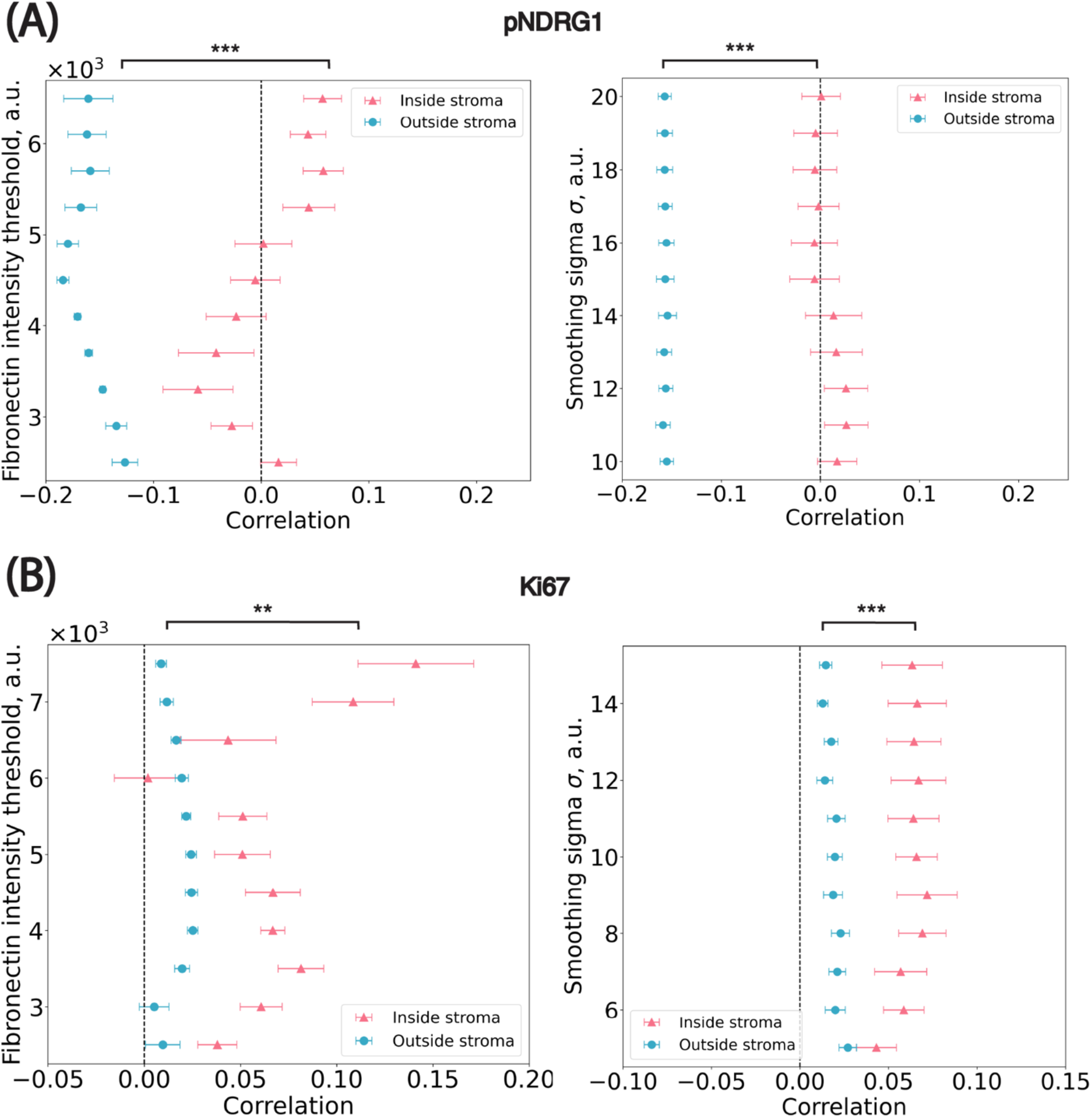
Sensitivity analysis on parameters used in the stroma annotation. **(A)** pNDRG1 dataset. ** *p* = 1.95 × 10^−3^; *** *p* = 9.77 × 10^−4^ **(B)** Ki67 dataset. *** *p* = 9.77 × 10^−4^; *** *p* = 9.77 × 10^−4^ x-axis: the median Pearson correlation value from all images in the dataset. Blue circular dots represent cells outside of stromal regions i.e. marked by positive distance to their closest border. Pink triangle dots represent cells inside stromal regions i.e. marked by negative distance to their closest border. Significance levels: * *p*<0.05, ** *p*< 0.01, *** *p*<0.001.

## 6. Discussion

In this study we present an open-source image analysis pipeline for the robust and reproducible quantification of spatial biomarker distributions in stroma-rich tumors. By integrating QuPath (Bankhead et al. 2017), StarDist (Schmidt et al. 2018), and Python-based statistical analyses, our framework enables the extraction of biologically relevant spatial patterns from heterogeneous immunofluorescence imaging datasets. Notably, we found that the fibronectin-defined stromal border consistently aligned with spatial changes in biomarker expression. Specifically, in cancer cells positive for the phospho-NDRG1, the signal intensity peaked at the tumor-stroma interface and started to rapidly decline in the cells located farther from the stromal border. As for the cells positive for Ki67, the intensity was low inside the stroma and increased progressively with greater distance from the border until plateauing at approximately 300 *µ*m mark (**Figure 5**).

These patterns reveal a spatially constrained organization of ECM sensing and proliferative response programs shaped by the surrounding stroma. While dense stromal regions influence the activation of key intracellular signaling pathways and restrict drug availability, other components of the TME orchestrate a wide range of immune, metabolic, and transcriptional changes that contribute to the complexity of stroma-rich cancers. Further biological and biochemical studies are needed to uncover the underlying cues driving the distinct spatial distribution of these programs.

### Scalability and generalizability

Our pipeline is designed to process gigapixel-scale whole-slide images with batch-level reproducibility. While classification via QuPath’s machine learning tools still requires manual annotation, we show that statistical propagation of thresholds from a reference image can approximate the accuracy of trained classifiers. This approach significantly improves scalability for large datasets by reducing the need for manual curation, while also compensating for staining variability across slides. One key take home message is that the framework is not limited to pancreatic cancer mouse xenografts, but can be applied for the analysis of various tissues with high stromal content (e.g. breast, ovarian cancer tissues, fibrotic tissues, bone marrow, or adipose tissue) (Friščić and Hoffmann 2022). Stromal regions can be delineated using any appropriate marker (e.g. collagens, laminins), making the pipeline adaptable to a wide range of tumor types and other dense tissues. Similarly, classification can be tailored to different biomarkers and experimental goals using either supervised or threshold-based methods, depending on the availability of labeled training data. This flexibility makes the pipeline applicable across diverse contexts even beyond cancer biology, where spatial phenotyping is relevant.

### 6.1. Limitations and future work

Despite its strengths, the pipeline has several limitations. First, stromal region delineation is currently based on intensity thresholding, which, although validated via sensitivity analysis, remains susceptible to staining variability and human bias. Incorporating texture features or adopting deep learning-based segmentation methods may further improve robustness by being even more agnostic for human-chosen parameters. Second, while the statistical propagation method mitigates batch effects, it still depends on the initial selection of a reference threshold. We show that this dependency is dataset-specific, and recommend examining cell-level intensity distributions before choosing between machine learning or percentile-based classification strategies. Additionally, this work focuses exclusively on 2D spatial measurements. Although informative, 2D projections may obscure spatial interactions that unfold in three dimensions. Future extensions of the pipeline could integrate volumetric imaging techniques and 3D segmentation to enable more comprehensive modeling of tumor-stroma interactions in space.

### 6.2. Conclusion

This study demonstrates a reproducible and scalable pipeline for analyzing the spatial organization of biomarkers in stroma-rich tissues using images acquired from the chemotherapy-treated pancreatic cancer xenografts as an example. By leveraging user-friendly open-source software and emphasizing interpretable statistical modeling, the approach lowers the barrier to quantitative spatial analysis in experimental pathology. As large-scale multiplexed imaging becomes increasingly common, tools like this will be essential for integrating spatial context into the study of tumor biology and therapy response.

## 7. Figures

### Box 1

Algorithm for statistically propagating classification thresholds across images based on the underlying distribution of cell measurements extracted from QuPath. It adjusts classification thresholds for each image by leveraging the statistical properties of cell marker intensity distributions. Given a reference threshold in a chosen image, the method translates this threshold to other images by aligning percentiles within the fitted distribution of cellular measurements. This approach accounts for variations in staining intensity and imaging conditions, ensuring consistency in classification across images.

#### Batch-wise statistical propagation of classification thresholds

Given a set of images *I*_1_,*I*_2_, …,*I*_*N*_ and for a specific channel *c*, we retrieve the histogram of each image’s pixel intensity distributions using a fixed number of bins, .*N*_*b*_.

1. **Histogram binning:**

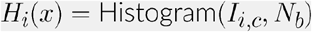

Where *H*_*i*_ (*x*) is the histogram of image *I*_*i*_ for channel *c*, and *x* the intensity levels.
2. **Least-square fit of a pool of non-negative distributions:** We fit a pool of non-negative distributions, *D*, on the intensity distributions:

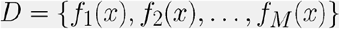

Here, *f*_*i*_(*x*) represents the *i*^th^ distribution function in the pool. The minimal pool of distributions contains common non-negative distributions: Log-normal, Wald, Burr, Beta and Gamma distributions. The best fit *f* ^*^(*x*) is retrieved using the least-squares method:

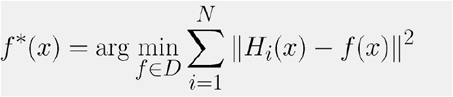
3. **Percentile mapping:** Given a pixel intensity threshold *t*_*i*_ in distribution *i*, its percentile is mapped to another distribution *j* using their cumulative distribution functions (CDFs):

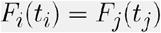

where *F*_*i*_ (*t*) and *F*_*j*_(*t*) are the CDFs of distributions *i* and *j*, respectively.
4. **Inverse probability calculation:** The value of *t*_*j*_ is then calculated using the inverse cumulative distribution function of distribution *j*:

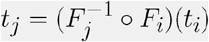

## 8. Data and code availability

All steps were performed on a workstation running Python 3.7 (Python Software Foundation, available at https://www.python.org) with standard scientific libraries (NumPy (Harris et al. 2020), SciPy (Virtanen et al. 2020), Pandas (McKinney 2010; The pandas development team 2024), Fitter, Matplotlib (Hunter 2007); a *uv* lock file is provided for dependencies). QuPath (v0.5+) was used for interactive segmentation, annotation, and feature export. Trained machine learning classifiers are available as .json files and can be loaded in QuPath to predict cell classes on new images or retrained on new annotations. The complete pipeline code is made open-source and is hosted on GitHub (https://github.com/HMS-IAC/stroma-spatial-analysis-web).

## 9. Author contributions

AAR, NK and SFN conceived the pipeline. AAR implemented the pipeline, trained ML models and performed data analysis. NK and KAC conceptualized and performed animal experiments. NK performed the immunostaining, acquired the images and annotated ROIs in whole-slides images. AAR and NK wrote the manuscript with input from SFN and TM. SFN and TM supervised the project. All authors contributed to the article and approved the submitted version.

## 10. Acknowledgements

This work was supported by Ludwig Center at Harvard (TM), NIH R01 CA258372 (TM), American Cancer Society grant RSG-19-0201-CSM (TM) and the Foundry Program at Harvard Medical School (SFN).

