## Supplementary tables and figures for "An image analysis pipeline to quantify the spatial distribution of cell markers in stroma-rich tumors"

**Paper website:** https://hms-iac.github.io/stroma-spatial-analysis-web

### Supplementary tables and figures

Numbers in tables were rounded up to the sixth significant decimal.

#### *AsPC NDRG1*

**Table S1:** Parameters of the fitted lognormal distribution for the ***FITC KER: Cytoplasm: Median*** intensity across all detected cells. The intensity threshold used for cell classification in each image is determined by translating the reference threshold percentile (marked with an *).

| Image | Location | Shape | Scale | Threshold |
|---|---|---|---|---|
| **#1** | –707.011083 | 0.619916 | 3138.058254 | 300* |
| **#2** | –723.643241 | 0.603754 | 2949.356883 | 251.28 |
| **#3** | –793.205413 | 0.618143 | 3229.167415 | 246.42 |
| **#4** | –308.258962 | 0.681428 | 1840.263228 | 219.30 |
| **#5** | –1129.335995 | 0.558484 | 4068.679791 | 331.98 |

**Table S2:** Parameters of the fitted lognormal distribution for the ***CY5 pNDRG1: Cell: Max*** intensity across all detected cells. The intensity threshold used for cell classification in each image is determined by translating the reference threshold percentile (marked with an *)

| Image | Location | Shape | Scale | Threshold |
|---|---|---|---|---|
| **#1** | 734.582782 | 1.361657 | 1067.563348 | 1200* |
| **#2** | 700.593146 | 1.238082 | 756.449602 | 1056.18 |
| **#3** | 703.018174 | 1.229991 | 828.940911 | 1094.61 |
| **#4** | 687.089880 | 1.151794 | 608.544201 | 988.61 |
| **#5** | 721.599776 | 1.516394 | 1077.707109 | 1149.14 |

**Table S3:** Parameters of the fitted lognormal distribution for the ***TRITC FN: Cell: Median*** intensity across all detected cells. The intensity threshold used for cell classification in each image is determined by translating the reference threshold percentile (marked with an *)

| Image | Location | Shape | Scale | Threshold |
|---|---|---|---|---|
| **#1** | 1127.574751 | 0.680330 | 1557.668026 | 5000* |
| **#2** | 1073.706558 | 0.741478 | 1482.000040 | 5072.28 |
| **#3** | 913.571101 | 0.573779 | 1331.199930 | 3783.08 |
| **#4** | 884.531585 | 0.670759 | 876.069113 | 3034.75 |
| **#5** | 1276.548976 | 0.727041 | 1644.706573 | 5629.18 |

#### ***AsPC Ki67***

**Table S4:** Parameters of the fitted lognormal distribution for the ***FITC KER: Cytoplasm: Median*** intensity across all detected cells. The intensity threshold used for cell classification in each image is determined by translating the reference threshold percentile (marked with an *)

| **Image** | **Location** | **Shape** | **Scale** | **Threshold** |
|---|---|---|---|---|
| **#1** | 288.717525 | 0.599332 | 1632.425859 | 650* |
| **#2** | -550.184474 | 0.564812 | 4726.485901 | 590.79 |
| **#3** | -9.668936 | 0.489416 | 2167.226010 | 622.80 |
| **#4** | -310.003757 | 0.416821 | 2126.460683 | 434.95 |
| **#5** | -340.941994 | 0.797041 | 8712.196935 | 831.45 |

**Table S5:** Parameters of the fitted lognormal distribution for the ***CY5 Ki67: Nucleus: Max*** intensity across all detected cells. The intensity threshold used for cell classification in each image is determined by translating the reference threshold percentile (marked with an *).

| **Image** | **Location** | **Shape** | **Scale** | **Threshold** |
|---|---|---|---|---|
| **#1** | 286.724866 | 1.801891 | 469.496465 | 950* |
| **#2** | 282.490001 | 1.643360 | 474.312730 | 932.51 |
| **#3** | 348.348119 | 1.593198 | 752.831402 | 1370.18 |
| **#4** | 241.491744 | 1.597186 | 255.996654 | 589.23 |
| **#5** | 260.713033 | 1.705415 | 707.838623 | 1242.37 |

**Table S6:** Parameters of the fitted lognormal distribution for the ***FITC FN: Cell: Median*** intensity across all detected cells. The intensity threshold used for cell classification in each image is determined by translating the reference threshold percentile (marked with an *).

| Image | Location | Shape | Scale | Threshold |
| --- | --- | --- | --- | --- |
| **#1** | 695.439669 | 0.477924 | 1499.172311 | 4000* |
| **#2** | 1053.805650 | 0.663097 | 1866.380304 | 6641.85 |
| **#3** | 1258.215495 | 0.603436 | 1219.408835 | 4566.16 |
| **#4** | 564.392175 | 0.634543 | 718.958016 | 2617.70 |
| **#5** | 805.830558 | 0.649840 | 1221.342242 | 4383.30 |

**Table S7:** Agreement percentages of single-threshold classifiers when compared to machine learning classification results for pNDRG1 images. The same thresholds from one image were applied to classify cells in other images. $\sigma$ is the sample standard deviation. All values are in percentage (%).

| **ML classifier** | **Classifier using thresholds from image number:** | | | | |
| --- | --- | --- | --- | --- | --- |
| | **#1** | **#2** | **#3** | **#4** | **#5** |
| 87.7 | 96.1 | 86.4 | 90.1 | 78.7 | 94.0 |
| | $\mu = 89.06$<br>$\sigma = 6.88$ | | | | |

**Table S8:** Agreement percentages of single-threshold classifiers when compared to machine learning classification results for Ki67 images. The same thresholds from one image were applied to classify cells in other images. σ is the sample standard deviation. All values are in percentage (%).

<table>
<tr><th>ML classifier</th><th colspan="5">Classifier using thresholds from image number:</th></tr>
<tr><td rowspan="3">86.3</td><td>#1</td><td>#2</td><td>#3</td><td>#4</td><td>#5</td></tr>
<tr><td>84.2</td><td>83.7</td><td>92.3</td><td>66.6</td><td>90.4</td></tr>
<tr><td colspan="5">$\mu = 83.4$<br>$\sigma = 10.1$</td></tr>
</table>

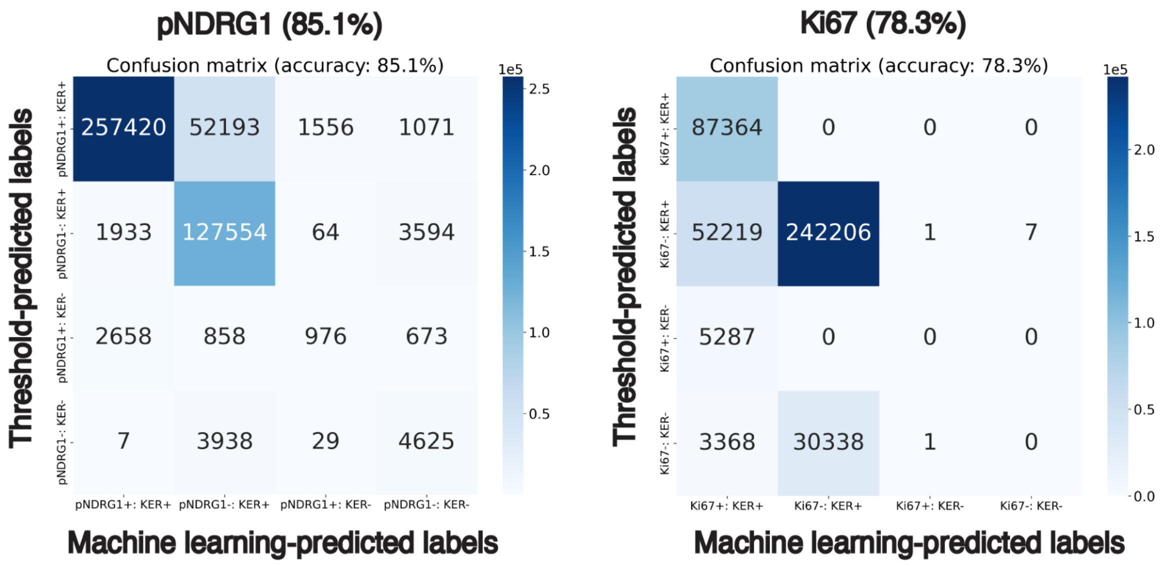


**Figure S1:** Comparison of machine learning-based and threshold-based classifiers across all possible class assignments from the following four classes: KER+, KER-, pNDRG1- or Ki67+, and pNDRG1- or Ki67-.

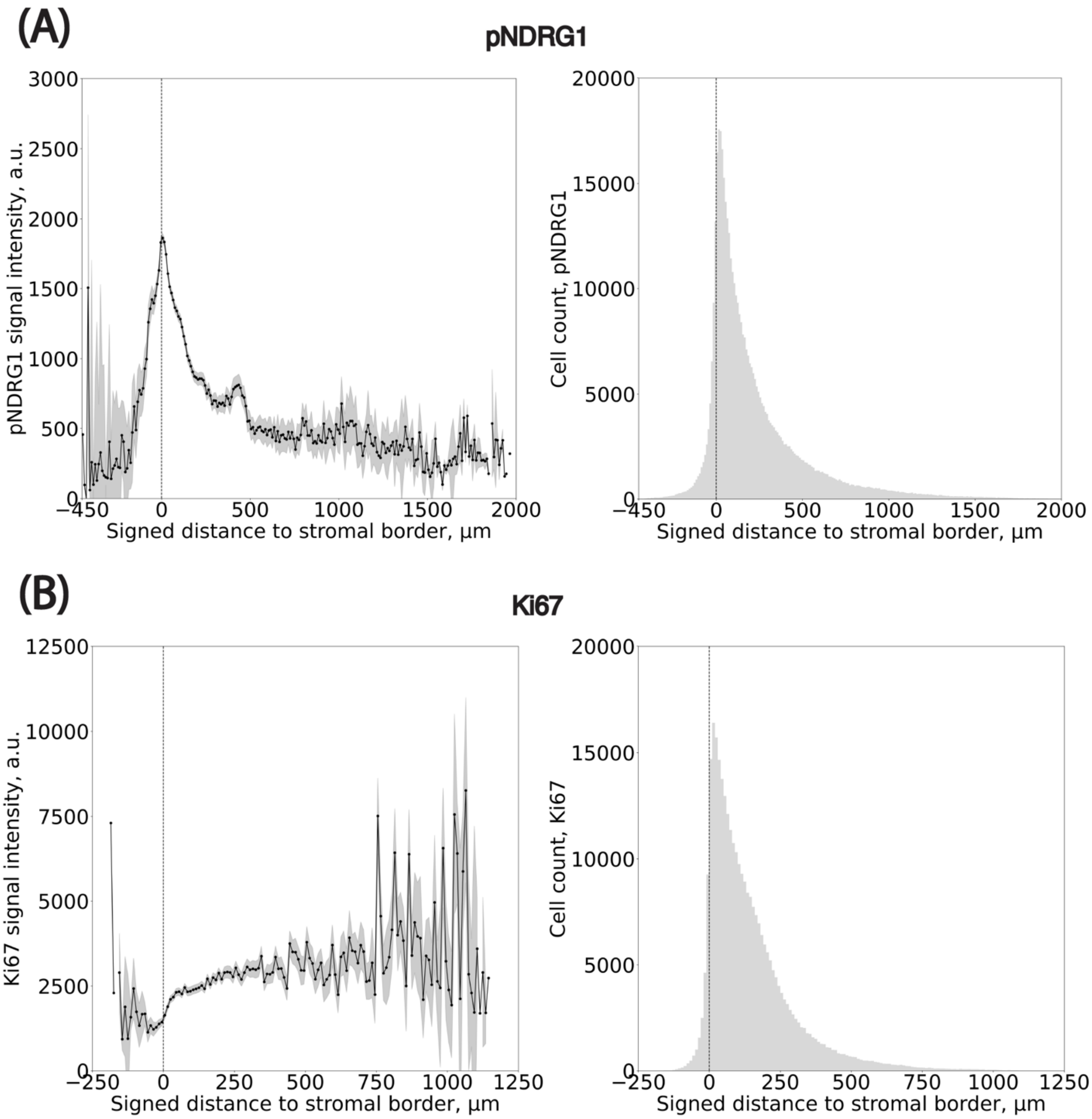


**Figure S2:** Full spatial distribution of double positive cells (cytokeratin positivity and pNDRG1 or Ki67) relative to the modelled stromal border.

**(A)** (left) Full spatial distribution of pNDRG1-positive cancer cells. Bin size: 10 $\mu$m. x axis is unclipped. Black dots represent the median value within a bin. Grey overlays indicate Standard Error of the Mean [SEM]. (right) Number of cells per bin. Same bin size. **(B)** Same plots repeated for Ki67-positive cancer cells. Bin size: 10 $\mu$m.
